# Negative effects characterization and comparative transcriptomics elucidation on the lag phase of an industrial *S. cerevisiae* under the corn stover hydrolysate stress

**DOI:** 10.1101/2020.03.16.994723

**Authors:** Xiaolin Kuang, Yaping Guo, Zhengyue Zhang, Xiangdong Hu, Xuebing Han, Yidan Ouyang, Difan Xiao, Qian Li, Hanyu Wang, Xi Li, Qiang Chen, Menggen Ma

**Author notes:** Xiaolin Kuang and Yaping Guo contributed equally to this work. Address correspondence to Menggen Ma,. Sichuan Society for Microbiology, Chengdu, Sichuan 611130, P. R. China.

## Abstract

During biofuels fermentation from pretreated lignocellulosic biomass, the strong toxicity of the lignocellulose hydrolysate is resulted from the synergistic effect of multiple lignocellulosic inhibitors, which far exceeds the sum of effects caused by every single inhibitor. Meanwhile, the synergistic effect is unclear and the underlying response mechanism of the industrial yeast towards the actual pretreated lignocellulose hydrolysate is still under exploration. Here, we employed an industrial *S. cerevisiae* for the transcriptomic analysis in two time points (early and late) of the lag phase under the corn stover hydrolysate stress. As investigation, the corn stover hydrolysate caused the accumulation of reactive oxygen species (ROS), damages of mitochondrial membrane and endoplasmic reticulum (ER) membrane in the industrial *S. cerevisiae* YBA_08 during the lag phase, especially these negative effects were more significant at the early lag phase. Based on the transcriptome profile, the industrial *S. cerevisiae* YBA_08 might recruit stress-related transcription factors (*MSN4*, *STE12*, *SFL1*, *CIN5*, *COM2*, *MIG3*, etc.) through the mitogen-activated protein kinase (MAPK)-signaling pathway to induce a transient G1/G2 arrest, and to activate defense bioprocesses like protectants metabolism, sulfur metabolism, glutaredoxin system, thioredoxin system, heat shock proteins chaperone and oxidoreductase detoxification, resisting those compounded stresses including oxidative stress, osmotic stress and structural stress. Surprisingly, this defense system might be accompanied with the transient repression of several bioprocesses like fatty acid metabolism, purine *de novo* biosynthesis and ergosterol biosynthesis.

**Importance:** This research systematically demonstrated the lag phase response of an industrial yeast to the lignocellulosic hydrolysate in transcriptional level, providing a molecular fundament for understanding the synergistic effect of various lignocellulosic inhibitors and the regulatory mechanism of tolerance for industrial yeasts under this stress.

## Introduction

As a potential biopolymer for renewable biofuel production and block chemical building, lignocellulosic biomass has been extensively used (1, 2). Indeed, more than 60% of total biomass exist in the form of agricultural residues, such as wheat straw, rice straw and corn stover, all of which are composed of cellulose, hemicellulose and lignin (3, 4). However, apart from fermentable sugars, excessive unwilling inhibitors like aldehyde, phenolic and aliphatic acid are released through the pretreatment process, which negatively affect biocatalysts growth and fermentation (5, 6). Accounting for economic efficiency, the inhibitor-resistant biocatalyst engineered by genetic approaches can overcome the problem of the expensive cost for removing inhibitors (7, 8). Among several biocatalysts, *S. cerevisiae* exhibits the superiority in plastic genome and more resistance to lignocellulosic inhibitors than others like *Z. mobilis*. So it is regarded as one of the most valuable organisms to dissect negative physiological effects and the molecular tolerance mechanism under the lignocellulosic inhibitor stress (9, 10).

In last years, multitudinous investigations have been carried out and reveal that lignocellulosic inhibitors can cause *S. cerevisiae* metabolic disturbances, such as reactive oxygen species (ROS) accumulation and redox imbalance, and structural defects including plasma membrane instability, cell wall integrity loss and mitochondrial damages during biofuel fermentation (11). With the involvement of omics technologies in eliciting lignocellulosic inhibitor tolerance mechanism, significant advances have been made (12, 13). Stress-related biological processes and key genes capable of resisting lignocellulosic inhibitors have been revealed, providing ways for improving biofuel industrial fermentation through modifying yeast genotype. According to previous reports, they demonstrate that yeast can reconstitute metabolism pathways to resist furfural and 5-hydroxymethyl-2-furaldehyde (HMF) stress by initially enhancing mitogen-activated protein kinase (MAPK)-signaling pathway like filamentous growth pathway (FG), high osmolarity glycerol pathway (HOG) and cell wall integrity pathway (CWI), and subsequently activating transcription factor (TF)-encoding genes, such as *DIG1*, *MSN2*/*MSN4*, *SWI4* and *HSF1*, to regulate downstream stress-responsive metabolism pathways (14, 15). And the activated key bioprocesses include pentose phosphate (PPP) pathway (*ZWF1*, *GND1* and *TDH1*), proline biosynthesis (*PUT1* and *PUT2*) and multidrug efflux pump (*TPO1*, *TPO2* and *SNQ2*) (15, 16). As for response to acetic acid stress, genes involved in glycolysis pathway (*HXK2* and *PFK1*), ATP synthesis (*ATP1* and *ATP5*) and H^+^ transporters (*PMA1* and *PMP2*) are required for maintaining intracellular redox and ion homeostasis in *S. cerevisiae* (17). Upon mixture of furfural and acetic acid, biosynthesis of these protectants like trehalose and 4-aminobutyrate (GABA) is enhanced to provide additional defenses for *S. cerevisiae* against oxidative and redox stresses (18). Although the adaptive response mechanism towards single or multiple lignocellulose inhibitors, such as furfural, HMF and acetic acid, have been extensively demonstrated based on lab yeasts, the systematic analysis of industrial yeasts response to the lignocellulose hydrolysate generated by industrial pretreatments has been generally less approached due to the unclear synergistic effect of complex mixtures in the lignocellulose hydrolysate (19, 20). Therefore, it is essential and meaningful to aim to the industrial yeast to investigate the synergistic effect under the compounded lignocellulose hydrolysate stress, and to elicit the underlying genetic regulatory network and the genetic perturbation target.

In order to reveal valuable insights into the molecular response mechanism of industrial yeasts towards the lignocellulose hydrolysate, the lag phase transcriptomic profile of an industrial yeast *S. cerevisiae* YBA_08 was performed by RNA-seq in this study. These data will be available for understanding the molecular mechanism for resistance to the complex lignocellulose hydrolysate and obtaining resistant-yeasts engineered by reverse genetic methods.

## Results

### Negative effects characterization under the corn stover hydrolysate stress

#### Growth in YPH

During incubation in YPH medium *S. cerevisiae* YBA_08 distinctly exhibited a longer lag phase than being cultured in YPD medium, and ultimately could reach the maximal optical density, indicating that this corn stover hydrolysate negatively affected *S. cerevisiae* YBA_08 growth (Fig. 1).

**Fig. 1.**
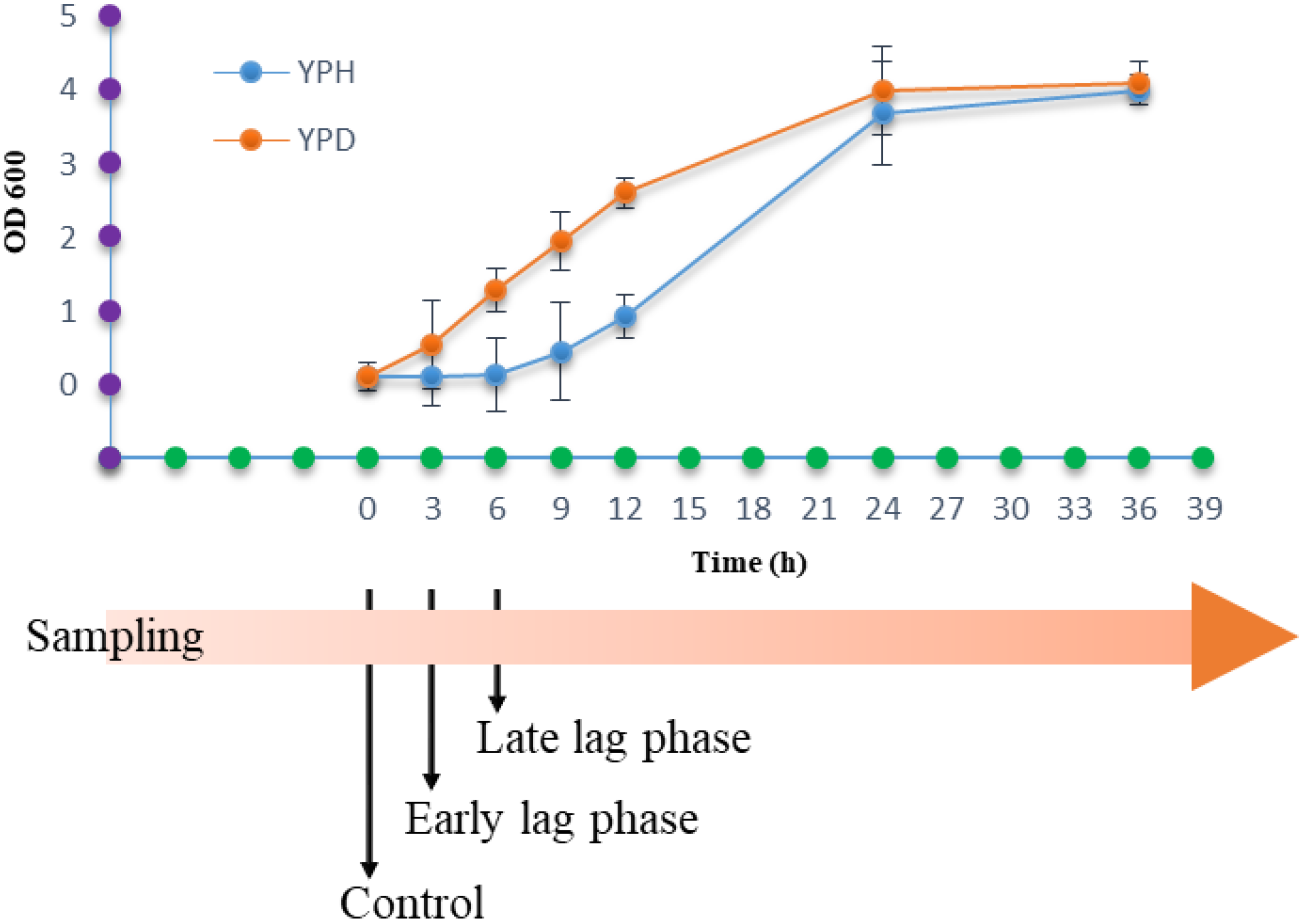
Growth of the industrial *S. cerevisiae* YBA_08 in YPH and the experimental set-up for transcriptome profile. During growing in YPH medium, the strain distinctly exhibited a longer lag phase than being cultured in YPD medium. The experimental set-up was displayed below. Samples for RNA-seq were prepared at different time points after incubation under YPH medium, including the early lag phase (3 h) and the late lag phase (6 h). Control samples were taken at 0 h.

#### ROS accumulation, mitochondrial and ER membrane damages

In ROS accumulation analysis (Fig. 2a), the industrial *S. cerevisiae* YBA_08 cells staining positive for ROS at 3 h (early) and at 6 h (late) were 87.4% and 54.1%, respectively, whereas the cells at 0 h (the control) was 13.9%. Comparing with the control, both of positive rates at 3 h and at 6 h exhibited significant differences (P < 0.01). After stained by MitoTracker Green FM, the normal mitochondria of the industrial *S. cerevisiae* YBA_08 showed typically tubular well-distributed under microscopy, and the damaged one displayed fragmented and aggregated. The damage rates detected at 0 h, at 3 h and at 6 h were 10.7%, 83.9% and 70.4%, respectively (Fig. 2b). Both of damage rates at 3 h and at 6 h exhibited significant differences (P < 0.01). As investigated, the rates of the abnormal ER with disorganized and diffuse membrane network at 0 h, at 3 h and at 6 h were 6.0%, 47.0% and 15.9%, respectively (Fig. 2c). Both of the damage rates at 3 h and at 6 h showed significant differences (P < 0.01). Overall, this corn stover hydrolysate could induce ROS accumulations, mitochondrial and ER membrane damages in the industrial *S. cerevisiae* YBA_08, and these negative effects were more serious at 3 h.

**Fig. 2.**
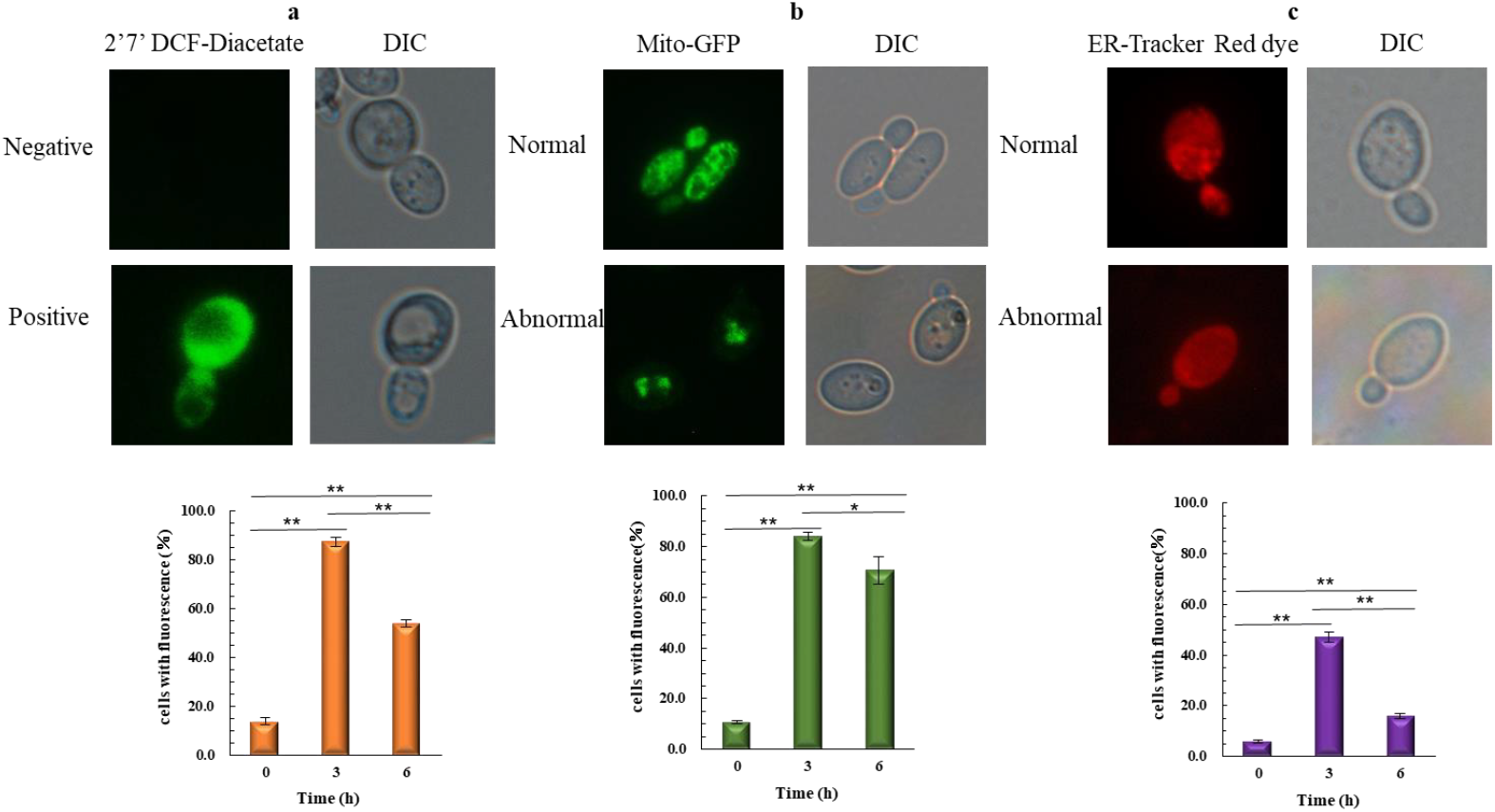
The corn stover hydrolysate induced ROS accumulation, mitochondria membrane damage and ER membrane damage. **a** The images of the industrial *S. cerevisiae* YBA_08 cells with a positive or negative ROS signal under fluorescence microscopy, and the ROS positive rates at 0 h (no inhibitors), at 3 h (early) and at 6 h (late). **b** The morphology of normal and abnormal mitochondria under fluorescence microscopy, and the quantification of cells with abnormal mitochondria at 0 h, at 3 h and at 6 h. **c** The morphology of normal and abnormal ER under fluorescence microscopy, and the quantification of cells with abnormal ER at 0 h, at 3 h and at 6 h. The experiments were carried out in triplicate. The asterisk (*) and (**) indicated the significant difference at P < 0.05 and at P < 0.01, respectively.

### Transcriptome differential expression analysis

To reveal the lag phase transcriptional response towards the lignocellulose hydrolysate, an industrial *S. cerevisiae* YBA_08 was examined under the corn stover hydrolysate (Fig. 1). All DEGs at early (3 h) and late (6 h) lag phases were both categorized to analyze temporal responses of DEGs after challenged with this stress. The DEGs analysis was conducted by comparing to the transcriptional profile under the control (0 h) (Fig. S1). Among the yeast genome, 799 transcripts were identified as significantly altered DEGs to early respond to this corn stover hydrolysate stress, including 404 up-regulated DEGs and 395 down-regulated DEGs (Fig. S1). Based on functional catalogues analysis through MIPS Functional Catalogue Database, 13 up-regulated catalogues (Fig. 3a) and 5 down-regulated catalogues (Fig. 3b) were characterized as key bioprocesses involved in coping with the toxic effect of this corn stover hydrolysate at the early lag phase. And these varied functional catalogues reflected synergistic effects among bioprocesses in resisting this stress. For example, under this stress, DEGs mapped to catalogues, such as stress response (MIPS ID:32.01), C-compound and carbohydrate metabolism (MIPS ID:01.05), amino acid metabolism (MIPS ID:01.01), detoxification (MIPS ID:32.07), C-compound and carbohydrate transport (MIPS ID:20.01.03) and protein folding and stabilization (MIPS ID:14.01) were significantly enhanced, and DEGs scored to catalogues including fatty acid metabolism (MIPS ID:01.06.05) and purine nucleotide/nucleoside/nucleobase anabolism (MIPS ID:01.03.01.03) were significantly repressed (Fig. 3 and Table S1). Additionally, the late response to this corn stover hydrolysate stress comprised 511 up-regulated DEGs and 355 down-regulated DEGs (Fig. S1). Most of up-regulated DEGs were enriched in functional catalogues such as ribosome biogenesis (MIPS ID:12.01), nucleic acid binding (MIPS ID:16.03) and translation (MIPS ID:12.04) (Fig. 3c and Table S2). And the majority of down-regulated DEGs were classified into transported compounds (substrates) (MIPS ID:20.01), cellular import (MIPS ID:20.09.18) and secondary metabolism (MIPS ID: 01.20) (Fig. 3d and Table S2). Overall, these data indicated that the industrial *S. cerevisiae* YBA_08 reprogrammed intracellular bioprocesses to adapt to this stress at the lag phase.

**Fig. 3.**
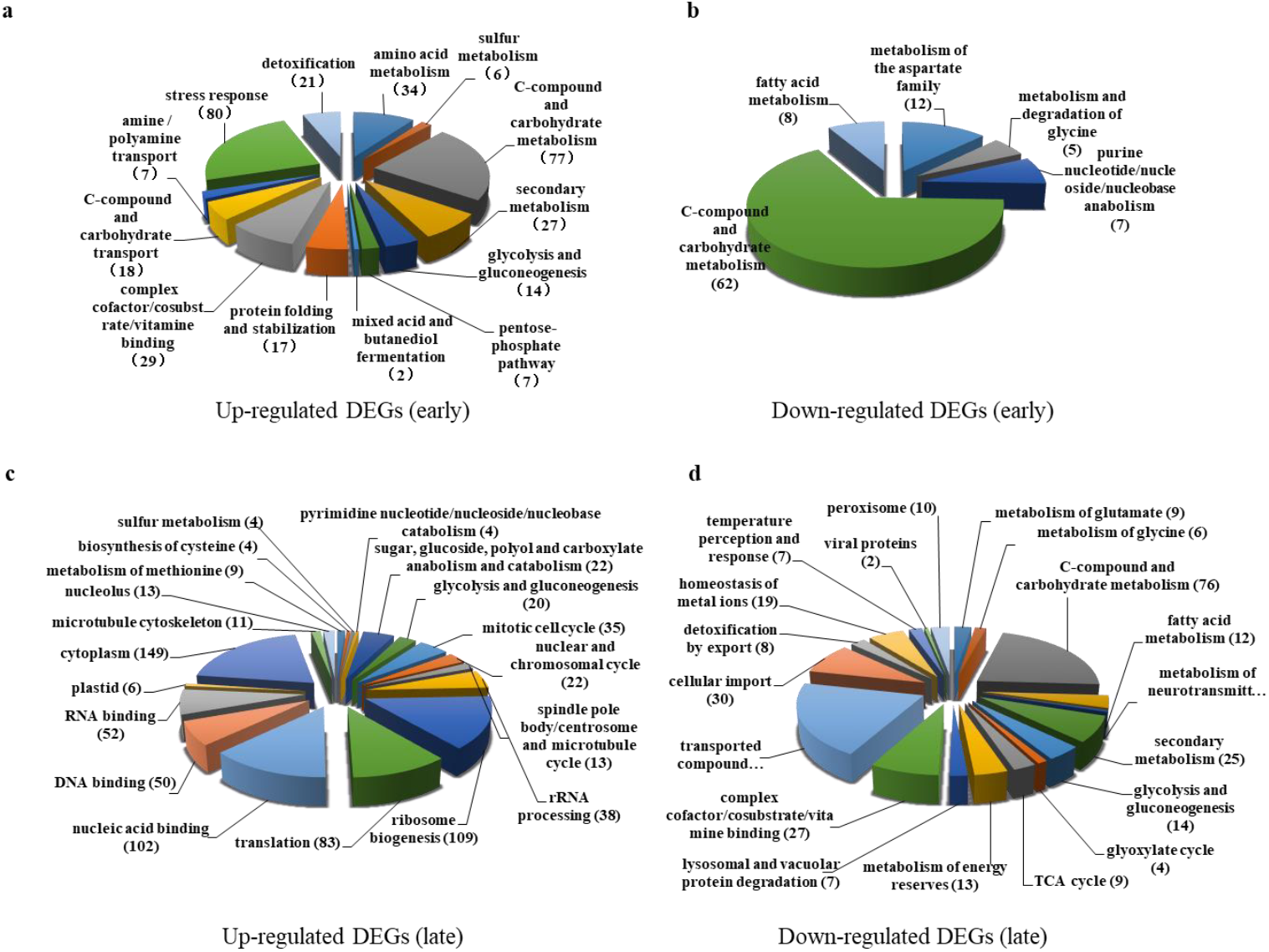
Functional catalogues analysis of early and late DEGs during lag phase identified from this transcriptome profile. **a** Functional catalogues of up-regulated early DEGs. **b** Functional catalogues of down-regulated early DEGs. **c** Functional catalogues of up-regulated late DEGs. **d** Functional catalogues of down-regulated late DEGs. The number followed behind each functional catalogue was the quantification of DEGs involved.

### Carbohydrate metabolism response

Many previous studies have demonstrated that monosaccharides like glucose, xylose and galactose could be released into lignocellulose hydrolysate after treatment by cellulose (21, 22). Under this corn stover hydrolysate, the early transcriptome profile of the industrial *S. cerevisiae* YBA_08 demonstrated that hexose transporter-encoding genes, such as *HXT1*, *HXT4*, *HXT7* and *GAL2*, were activated and genes involved in initiating glucose (*HXK2*), galactose (*GAL1*, *GAL7*, *GAL10* and *PGM2*) and xylose (*GRE3* and *XYL2*) to enter subsequent processes of glycolysis pathway were significantly enhanced (Table 1 and Fig. 4). Especially *GAL1*, *GAL2*, *GAL7* and *GAL10* showed high alterations with fold changes of 331.0–561.1 at the early lag phase, which were extremely higher than any other DEGs. Furthermore, genes in the process of converting glucose-6-P to glyceraldehyde-3P (*PGI1*, *YNR071C* and *FBA1*) and the process of converting glyceraldehyde-3P to pyruvate (*PGK1*, *YCR013C*, *ENO1* and *ERR3*) in glycolysis pathway were up-regulated (Table 1 and Fig. 4). Also, the key genes (*ZWF1*, *SLO1*, *SLO4*, *GND2*, *YBR116C* and *NQM1*) in PPP pathway were examined to respond positively under this stress, and these transcriptional abundances were actually high. Additionally, metabolisms of osmolytes like trehalose, glycogen and glycerol obviously displayed (Table 1 and Fig. 4) enhancement according to the up-regulation of genes in trehalose metabolism (*UGP1*, *TPS1*, *TPS2*, *TSL1*, *NTH1* and *NTH2*), in glycogen metabolism (*GSY2*, *GDB1* and *GPH1*), and in glycerol metabolism (*GPP1*, *GCY1*, *DAK1* and *DAK2*). Meanwhile, genes in glycerol channels (*ASK10*, *STL1* and *GUP1*) were activated under this stress (Table 1). Consequently, the majority of above genes were not consistently activated, and expression abundances returned to normal at the late lag phase except genes in glycolysis (*GAL1*, *GAL2*, *GAL7*, *GAL10*, *HXK2*, *PGI1*, *FBA1*, *PGK1*, *YCR013C* and *ENO1*), and genes in glycerol metabolism (*GPP1*, *GCY1*, *STL1* and *GUP2*). The transcription level of *GAL1*, *GAL2*, *GAL7* and *GAL10* were extremely high at the late lag phase.

**Fig. 4.**
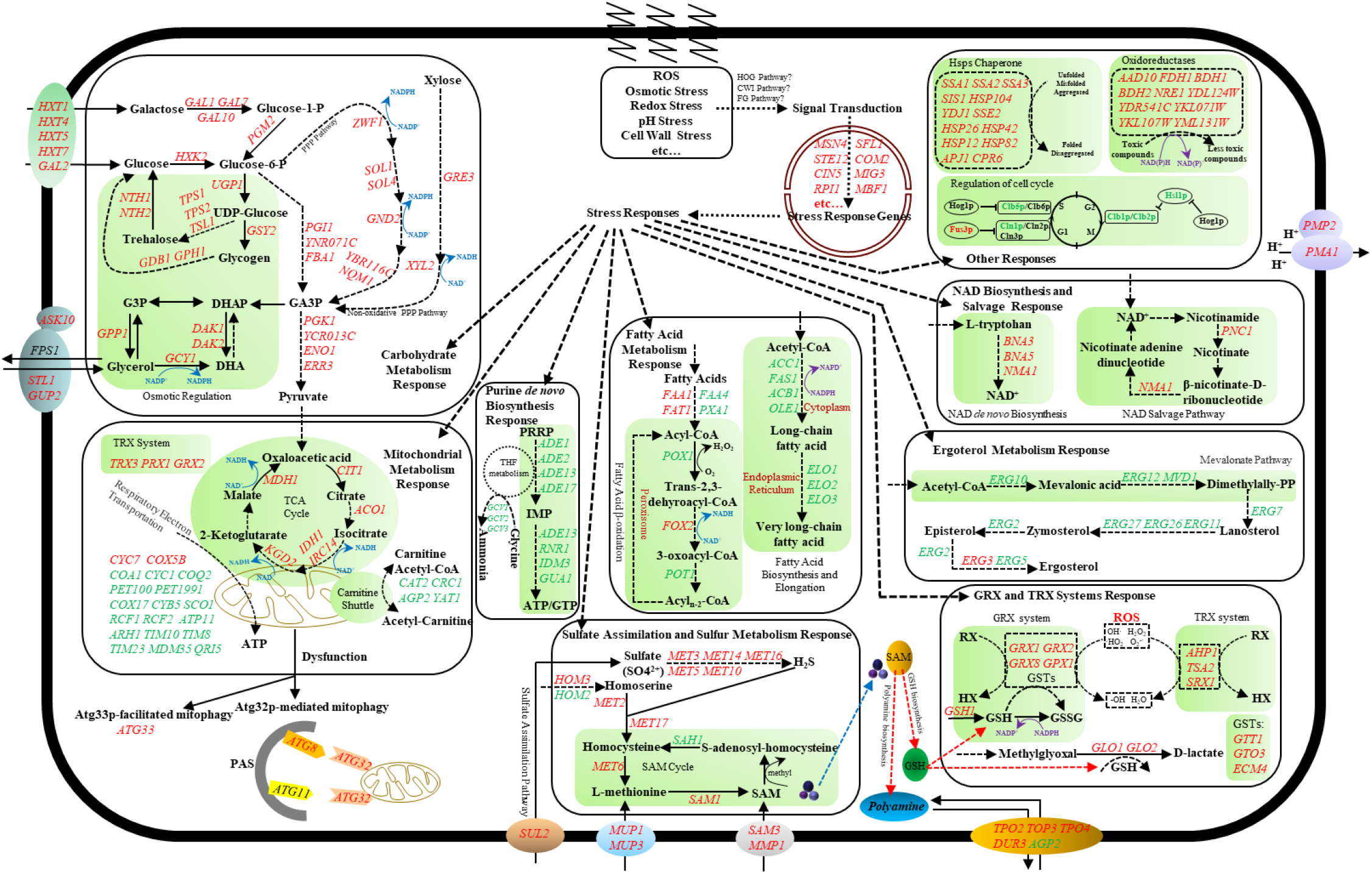
The early lag phase transcriptional response of the industrial *S. cerevisiae* YBA_08 to the corn stover hydrolysate stress. After exposure to the corn stover hydrolysate, the induction of stress including oxidative stress, osmotic stress and structural stress, recruited stress-responsive genes to defense through MAPK signaling pathways including FG pathway, HOG pathway and CWI pathway. The stress-responsive genes were involved in carbohydrate metabolism, mitochondrial metabolism, fatty acid metabolism, purine *de novo* biosynthesis, sulfur metabolism, ergosterol metabolism, NAD biosynthesis and salvage, GRX and TRX systems, regulation of cell cycle, Hsps chaperone and oxidoreductases. Genes with red color indicated significantly up-regulation (fold change≽2); Genes with purple color displayed slightly up-regulation (1.4≼fold change<2); Genes with green color indicated significantly down-regulation (fold change≽2); Genes with blue color displayed slightly down-regulation (1.4≼fold change<2). MAPK: mitogen-activated protein kinases; FG: filamentous growth; HOG: high osmolarity glycerol; CWI: cell wall integrity; ROS: reactive oxygen species; HSPs: heat shock proteins; TCA: tricarboxylic acid: THF: tetrahydrofolate metabolism; IMP: inosine monophosphate; PRPP: 5-phospho-α-D-ribose 1-diphosphate; SAM: S-adenosylmethionine; GSH: glutathione; GSSG: oxidized glutathione; GRX: glutaredoxin; TRX: thioredoxin; GPXs: glutathione peroxidases; GSTs: glutaredoxin transferases; PAS: phagophore assembly site; RX: xenobiotics; HX: reduced xenobiotics, PPP: pentose phosphate; G3P: glycerol-3P GA3P: glyceraldehyde-3P; DHAP: dihydroxyacetone phosphate; DHA: dihydroxyacetone.

**Table 1.** Comparative expression fold changes of carbohydrate metabolism-related genes for the industrial *S. cerevisiae* YBA_08 response to this corn stover hydrolysate at early and late lag phases.

| Gene | Description | Fold change |  | Gene | Description | Fold change |  |
| --- | --- | --- | --- | --- | --- | --- | --- |
|  |  | Early lag phase | Late lag phase |  |  | Early lag phase | Late lag phase |
| Glycolysis |  |  |  |  |  |  |  |
| HXT1 | Low-affinity glucose transporter | 2.8 | - | PGI1 | Glycolytic enzyme phosphoglucose isomerase | 2.0 | 2.3 |
| HXT4 | High-affinity glucose transporter | 4.7 | - | YNR071C | Putative aldose 1-epimerase | 4.8 | - |
| HXT5 | Hexose transporter | 1.6 | -4.0 | FBA1 | Fructose 1,6-bisphosphate aldolase | 1.6 | 2.5 |
| HXT7 | High-affinity glucose transporter | 25.1 |  | PGK1 | 3-phosphoglycerate kinase | 1.7 | 2.0 |
| GAL1 | Galactokinase | 331.0 | 521.2 | YCR013C | Dubious open reading frame | 1.9 | 2.6 |
| GAL2 | Galactose permease | 561.1 | 816.5 | ENO1 | Enolase I, a phosphopyruvate hydratase | 2.3 | 3.1 |
| GAL7 | Galactose-1-phosphate uridyl transferase | 542.2 | 955.4 | ERR3 | Enolase, a phosphopyruvate hydratase | 3.4 | - |
| GAL10 | UDP-glucose-4-epimerase | 472.3 | 755.7 | GRE3 | Aldose reductase | 5.2 | - |
| HXK2 | Hexokinase isoenzyme 2 | 2.7 | 8.3 | XYL2 | Xylitol dehydrogenase | 2.4 | -2.6 |
| PGM2 | Phosphoglucomutase | 3.2 | - |  |  |  |  |
| PPP pathway |  |  |  |  |  |  |  |
| ZWF1 | Glucose-6-phosphate dehydrogenase | 1.7 | - | GND2 | 6-phosphogluconate dehydrogenase | 8.1 | - |
| SOL1 | homologous to Sol3p and Sol4p | 2.3 | - | YBR116C | Dubious open reading frame | 13.0 | - |
| SOL4 | 6-phosphogluconolactonase | 4.2 | -2.9 | NQM1 | Transaldolase of unknown function | 5.4 | - |
| Trehalose and Glycogen metabolism |  |  |  |  |  |  |  |
| UGP1 | UDP-glucose pyrophosphorylase | 1.6 | - | GDB1 | Glycogen debranching enzyme | 2.1 | -2.4 |
| TPS1 | Synthase subunit of trehalose-6-P synthase/phosphatase complex | 2.9 | - | GPH1 | Glycogen phosphorylase | 2.6 | - |
| TPS2 | Phosphatase subunit of the trehalose-6-P synthase/phosphatase complex | 2.1 | - | NTH1 | Neutral trehalase | 1.5 | - |
| GSY2 | Glycogen synthase | 1.7 | -2.1 | NTH2 | Putative neutral trehalase | 1.8 | - |
| Glycerol metabolism |  |  |  |  |  |  |  |
| GPP1 | Constitutively expressed DL-glycerol-3-phosphate phosphatase | 3.9 | 4.0 | AQY3 | Putative channel-like protein; similar to Fps1p | 5.7 | - |
| GCY1 | Glycerol dehydrogenase | 5.0 | 5.0 | STL1 | Glycerol proton symporter of the plasma membrane | 4.9 | 4.5 |
| DAK1 | Dihydroxyacetone kinase | 6.9 | - | GUP2 | Possible role in proton symport of glycerol | 1.4 | 3.5 |
| DAK2 | Dihydroxyacetone kinase | 3.7 | - | AQY2 | Water channel that mediates water transport across cell membranes | -1.4 | 3.1 |
| ASK10 | Regulator of the Fps1p glycerol channel | 2.4 | - |  |  |  |  |

"-": the differential expression was detected to be normal.

### Fatty acid and Ergosterol metabolism response

Reportedly, fatty acid biosynthesis, elongation and β-oxidation were occurred in cytoplasm, on ER and in peroxisome, respectively (23, 24). This transcriptome profile of the industrial *S. cerevisiae* YBA_08 for the early response indicated that most of genes related to fatty acid metabolism mentioned above were observed to be repressed to some extent (Table 2 and Fig. 4). In fatty acid biosynthesis and elongation, seven essential genes (*ACC1*, *FAS1*, *ACB1*, *OLE1*, *ELO1*, *ELO2* and *ELO3*) in the process from acetyl-CoA to very long-chain fatty acid were down-regulated. Moreover, several essential genes necessary for fatty acid β-oxidation, such as *FAA4*, *PXA1*, *POX1* and *POT1*, were repressed except *FAA1*, *FAT1* and *FOX2*. Notably, under this stress early expressions of most of genes (*ERG10*, *ERG12*, *MVD1*, *ERG7*, *ERG11*, *ERG26*, *ERG27*, *ERG2* and *ERG5*) in ergosterol metabolism were negatively regulated. Together with the late transcriptomic analysis, genes (*FAA4*, *PXA1*, *POX1*, *POT1*, *ACC1* and *FAS1*) in fatty acid metabolism showed a consistent down-regulation over all sampling points.

**Table 2.** Comparative expression fold changes of genes involved in fatty acid metabolism, ergosterol metabolism, NAD biosynthesis, NAD salvage and regulation of cell cycle for the industrial *S. cerevisiae* YBA_08 response to this corn stover hydrolysate at early and late lag phases.

| Gene | Description | Fold change |  | Gene | Description | Fold change |  |
| --- | --- | --- | --- | --- | --- | --- | --- |
|  |  | Early lag phase | Late lag phase |  |  | Early lag phase | Late lag phase |
| Fatty acid metabolism |  |  |  |  |  |  |  |
| Fatty acid β-oxidation |  |  |  |  |  |  |  |
| FAA1 | Long chain fatty acyl-CoA synthetase | 2.3 | - | POX1 | Fatty-acyl coenzyme A oxidase | -3.2 | -2.3 |
| FAT1 | Very long chain fatty acyl-CoA synthetase and fatty acid transporter | 2.5 | - | FOX2 | 3-hydroxyacyl-CoA dehydrogenase and enoyl-CoA hydratase | 1.7 | - |
| FAA4 | Long chain fatty acyl-CoA synthetase | -3.0 | -2.0 | POT1 | 3-ketoacyl-CoA thiolase | -2.0 | -3.5 |
| PXA1 | Subunit of heterodimeric peroxisomal ABC transport complex | -1.9 | -5.1 | IDP3 | Peroxisomal NADP-dependent isocitrate dehydrogenase | -1.8 | -2.4 |
| Fatty acid biosynthesis and elongation |  |  |  |  |  |  |  |
| ACC1 | Acetyl-CoA carboxylase | -4.4 | -2.1 | ELO1 | Elongase I | -2.2 | - |
| FAS1 | Beta subunit of fatty acid synthetase | -5.5 | -2.2 | ELO2 | Fatty acid elongase | -2.4 | 4.0 |
| ACB1 | Acyl-CoA-binding protein | -2.0 | - | ELO3 | Elongase | -1.8 | 3.2 |
| OLE1 | Delta(9) fatty acid desaturase | -2.43 | -2.19 |  |  |  |  |
| Ergosterol metabolism |  |  |  |  |  |  |  |
| ERG10 | Acetyl-CoA C-acetyltransferase | -2.6 | - | ERG26 | C-3 sterol dehydrogenase | -2.2 | - |
| ERG12 | Mevalonate kinase | -2.4 | - | ERG27 | 3-keto sterol reductase | -2.1 | - |
| MVD1 | Mevalonate pyrophosphate decarboxylase | -3.3 | - | ERG2 | C-8 sterol isomerase | -2.0 | - |
| ERG7 | Lanosterol synthase | -1.9 | - | ERG3 | C-5 sterol desaturase | 1.8 | 12.6 |
| ERG11 | Lanosterol 14-α-demethylase | -1.5 | 2.1 | ERG5 | C-22 sterol desaturase | -1.8 | 2.8 |
| NAD biosynthesis and salvage |  |  |  |  |  |  |  |
| BNA3 | Kynurenine aminotransferase | 5.1 | 3.5 | NMA1 | Nicotinic acid mononucleotide adenyllyltransferase | 6.3 | 4.1 |
| BNA5 | Kynureninase | 1.7 | 2.0 | PNC1 | Nicotinamidase | 3.8 | - |
| Regulation of cell cycle |  |  |  |  |  |  |  |
| CLB5 | B-type cyclin | -2.5 | 2.0 | HSL1 | Nim1p-related protein kinase | -4.1 | - |
| CLB6 | B-type cyclin | - | 3.5 | SWE1 | Protein kinase that regulates the G2/M transition | -3.0 | - |
| CLN2 | G1 cyclin | -2.1 | 3.2 | CLB1 | B-type cyclin | -5.9 | 3.3 |
| FUS3 | Mitogen-activated serine/threonine protein kinase | 2.2 | - | CLB2 | B-type cyclin | -4.5 | 2.4 |

"-", the differential expression was detected to be normal.

### Mitochondrial metabolism response

At the early lag phase, data from RNA-seq revealed that great changes in mitochondrial metabolism (Table 3 and Fig. 4) of the industrial *S. cerevisiae* YBA_08 were examined out under this corn stover hydrolysate stress. Those changed processes comprised down-regulated genes essential for carnitine shuttle and for respiratory electron transportation, and up-regulated genes concerning TCA cycle, thioredoxin (TRX) system and mitophagy. For instant, genes (*CAT2*, *CRC1*, *AGP2* and *YAT1*) in carnitine shuttle and genes (*CBP3*, *COA1*, *COQ2*, *PET191*, *COX17*, *CYB5* and *SCO1*) in respiratory electron transportation showed varied degree of repression. And expression abundances of genes (*CIT1*, *ACO1*, *IDH1*, *IRC14*, *KGD2* and *MDH1*) related to generation of cofactor NADH in TCA cycle were extremely high. In the TRX system, two genes (*TRX3* and *PRX1*) responsible for scavenging ROS and xenobiotics were positively enhanced. Surprisingly, essential genes (*ATG8*, *ATG32* and *ATG33*) in Atg32p/Atg33p-mediated mitophagy were slightly up-regulated. Meanwhile, the early transcriptional profile also revealed that genes (*ARH1*, *TIM10*, *TIM8*, *TIM23*, *MDM35* and *QRI5*) in mitochondrial membrane and genes (*GCV1*, *GCV2*, *GCV3* and *SHM2*) in mitochondrial glycine metabolism were both negatively regulated (Table S3). Ultimately, most of early response genes in mitochondrial metabolism returned to normal after the temporal up-regulation according to the late transcriptional data, except that genes in carnitine shuttle were still negatively regulated (Table 3 and Table S3).

**Table 3.** Comparative expression fold changes of mitochondrial metabolism-related and transporters-encoding genes for the industrial *S. cerevisiae* YBA_08 response to this corn stover hydrolysate at early and late lag phases.

| Gene | Description | Fold change |  | Gene | Description | Fold change |  |
| --- | --- | --- | --- | --- | --- | --- | --- |
|  |  | Early lag phase | Late lag phase |  |  | Early lag phase | Late lag phase |
| Mitochondrial metabolism |  |  |  |  |  |  |  |
| TCA cycle |  |  |  |  |  |  |  |
| CIT1 | Citrate synthase | 2.3 | - | IRC14 | Dubious open reading frame | 2.2 | - |
| ACO1 | Aconitase | 2.1 | - | KGD2 | Dihydrolipoyl transsuccinylase | 2.1 | - |
| IDH1 | Subunit of mitochondrial NAD(+)-dependent isocitrate dehydrogenase | 2.4 | - | MDH1 | Mitochondrial malate dehydrogenase | 3.6 | - |
| Respiratory electron transportation |  |  |  |  |  |  |  |
| CYC7 | Cytochrome c isoform 2 | 2.8 | - | RCF2 | Cytochrome c oxidase subunit | 2.0 | - |
| COX5B | Subunit Vb of cytochrome c oxidase | 6.0 | - | PET100 | Chaperone that facilitates the assembly of cytochrome c oxidase | -1.6 | -2.1 |
| COX19 | Protein required for cytochrome c oxidase assembly | 1.8 | - | PET191 | Protein required for assembly of cytochrome c oxidase | -2.0 | -2.0 |
| CYC1 | Cytochrome c, isoform 1 | -1.9 | - | COX17 | Copper metallochaperone that transfers copper to Sco1p and Cox11p | -2.6 | - |
| CBP3 | Mitochondrial protein required for assembly of cytochrome bc1 complex | -2.1 | - | CYB5 | Cytochrome b5 | -6.8 | - |
| COA1 | Mitochondrial inner membrane protein | -2.2 | -2.1 | SCO1 | Copper-binding protein of mitochondrial inner membrane | -2.6 | -2.3 |
| COQ2 | Para hydroxybenzoate polyprenyl transferase | -3.8 | - | ATP11 | Molecular chaperone | -2.5 | - |
| RCF1 | Cytochrome c oxidase subunit | -1.9 | -2.2 | FMC1 | Mitochondrial matrix protein | -2.4 | - |
| Carnitine shuttle and TRX system |  |  |  |  |  |  |  |
| CAT2 | Carnitine acetyl-CoA transferase | -1.7 | -6.0 | AGP2 | Plasma membrane regulator of polyamine and carnitine transport | -1.6 | -3.1 |
| CRC1 | Mitochondrial inner membrane carnitine transporter | -2.5 | -4.4 | YAT1 | Outer mitochondrial carnitine acetyltransferase | -6.4 | -7.8 |
| TRX3 | Mitochondrial thioredoxin | 2.1 | - | PRX1 | Mitochondrial peroxiredoxin with thioredoxin peroxidase activity | 1.6 | - |
| Mitophagy |  |  |  |  |  |  |  |
| ATG8 | Component of autophagosomes and Cvt vesicles | 1.5 | -2.7 | ATG33 | Mitochondrial mitophagy-specific protein | 1.9 | -2.5 |
| ATG32 | Mitochondrial outer membrane protein required to initiate mitophagy | 1.4 | - | ATG31 | Autophagy-specific protein required for autophagosome formation | 1.6 | - |
| Polyamine transportation, Ion transporters and Others |  |  |  |  |  |  |  |
| TPO2 | Polyamine transporter of the major facilitator superfamily | 4.5 | 3.2 | HSP30 | Negative regulator of the H(+)-ATPase Pma1p | 7.8 | - |
| TPO3 | Polyamine transporter of the major facilitator superfamily | 2.9 | - | CTR1 | High-affinity copper transporter of plasma membrane | -164.4 | -115.8 |
| TPO4 | Polyamine transporter of the major facilitator superfamily | 2.4 | 1.9 | CTR3 | High-affinity copper transporter of the plasma membrane | -51.0 | -46.1 |
| DUR3 | Plasma membrane transporter for both urea and polyamines | 1.6 | - | QDR2 | Plasma membrane transporter of the major facilitator superfamily | 3.3 | 2.9 |
| AGP2 | Plasma membrane regulator of polyamine and carnitine transport | -1.6 | -3.1 | AQR1 | Plasma membrane transporter of the major facilitator superfamily | 5.5 | 3.0 |
| PMA1 | Plasma membrane P2-type H+-ATPase | 3.8 | 4.5 | ERC1 | Member of the multi-drug and toxin extrusion (MATE) family | 4.0 | 3.7 |
| PMP2 | Proteolipid associated with plasma membrane H(+)-ATPase (Pma1p) | 6.2 | 2.3 | GAP1 | General amino acid permease | -1.7 | -2.1 |

“-”, the differential expression was detected to be normal.

### Sulfate assimilation and Sulfur metabolism response

From RNA-seq data, it indicated that at the early lag phase (Table 4) a sulfate permease-encoding gene *SUL2*, most of genes in sulfate assimilation (*MET3*, *MET14*, *MET16*, *MET5* and *MET10*) and genes in sulfur metabolism (*HOM3*, *MET2*, *MET17*, *MET6* and *SAM1*) were highly activated. At the late lag phase (Table 4), most of genes were consistently activated except *SUL2*, *MTE14* and *MET6*. In addition, transcriptional levels of genes encoding for S-adenosylmethionine (SAM) permeases (*SAM3* and *MMP1*) and for methionine permeases (*MUP1* and *MUP3*), were high at the early lag phase and returned to normal at the late lag phase except *MUP3*.

**Table 4.** Comparative expression fold changes of genes involved in sulfur metabolism, purine metabolism, GRX system and TRX system for the industrial *S. cerevisiae* YBA_08 response to this corn stover hydrolysate at early and late lag phases.

| Gene | Description | Fold change |  | Gene | Description | Fold change |  |
| --- | --- | --- | --- | --- | --- | --- | --- |
|  |  | Early lag phase | Late lag phase |  |  | Early lag phase | Late lag phase |
| Sulfur metabolism |  |  |  |  |  |  |  |
| Sulfate assimilation |  |  |  |  |  |  |  |
| *SUL2* | High affinity sulfate permease | 6.7 | - | *MET10* | Subunit alpha of assimilatory sulfite reductase | 3.1 | 1.9 |
| *MET3* | ATP sulfurylase | 3.4 | 3.3 | *MET14* | Adenylylsulfate kinase | 2.3 | - |
| *MET5* | Sulfite reductase beta subunit | 4.8 | 2.5 | *MET16* | 3'-phosphoadenylylsulfate reductase | 4.9 | 2.6 |
| Homocysteine biosynthesis and SAM cycle |  |  |  |  |  |  |  |
| *HOM2* | Aspartic beta semi-aldehyde dehydrogenase | -1.5 | - | *SAH1* | S-adenosyl-L-homocysteine hydrolase | -2.4 | - |
| *HOM3* | Aspartate kinase (L-aspartate 4-P-transferase) | 3.6 | 4.0 | *SAM3* | High-affinity S-adenosylmethionine permease | 1.6 | - |
| *MET2* | L-homoserine-O-acetyltransferase | 4.8 | 2.0 | *MMP1* | High-affinity S-methylmethionine permease | 2.2 | - |
| *MET17* | O-acetyl homoserine-O-acetyl serine sulphydrylase | 9.1 | 4.2 | *MUP1* | High affinity methionine permease | 4.5 | - |
| *MET6* | Cobalamin-independent methionine synthase | 2.9 | - | *MUP3* | Low affinity methionine permease | 10.9 | 1.9 |
| *SAM1* | S-adenosylmethionine synthetase | 3.2 | 3.6 | *SAM4* | S-adenosylmethionine-homocysteine methyltransferase | -2.1 | 1.9 |
| Purine metabolism |  |  |  |  |  |  |  |
| *ADE1* | N-succinyl-5-aminoimidazole-4-carboxamide ribotide synthetase | -2.9 | -2.0 | *RNR1* | Major isoform of large subunit of ribonucleotide-diphosphate reductase | -2.4 | 5.4 |
| *ADE2* | Phosphoribosylaminoimidazole carboxylase | -2.1 | - | *RNR3* | Minor isoform of large subunit of ribonucleotide-diphosphate reductase | 2.3 | - |
| *ADE13* | Adenylosuccinate lyase | -2.6 | - | *YNK1* | Nucleoside diphosphate kinase | 2.1 | - |
| *ADE16* | Enzyme of 'de novo' purine biosynthesis | 1.7 | - | *IMD3* | Inosine monophosphate dehydrogenase | -2.0 | 2.7 |
| *ADE17* | Enzyme of 'de novo' purine biosynthesis | -3.8 | -2.6 | *GUA1* | GMP synthase | -1.7 | - |
| *GUD1* | Guanine deaminase | 2.4 | - |  |  |  |  |
| GRX and TRX systems |  |  |  |  |  |  |  |
| *GSH1* | Gamma glutamylcysteine synthetase | 1.6 | - | *GTT1* | ER associated glutathione S-transferase | 2.2 | -2.0 |
| *GRX1* | Glutathione-dependent disulfide oxidoreductase | 1.7 | -2.6 | *GLO1* | Monomeric glyoxalase I | 5.2 | - |
| *GRX2* | Cytoplasmic glutaredoxin | 1.9 | - | *GLO2* | Cytoplasmic glyoxalase II | 1.4 | - |
| *GRX8* | Glutaredoxin | 3.9 | 3.5 | *AHP1* | Thiol-specific peroxiredoxin | 3.3 | - |
| *GPX1* | Phospholipid hydroperoxide glutathione peroxidase | 4.8 | - | *TSA2* | Stress inducible cytoplasmic thioredoxin peroxidase | 5.4 | - |
| *GTO3* | Omega class glutathione transferase | 1.6 | -2.5 | *SRX1* | Sulfiredoxin | 2.2 | - |
| *ECM4* | S-glutathionyl-(chloro)hydroquinone reductase (GS-HQR) | 1.9 | -4.9 | *CTT1* | Cytosolic catalase T | 14.3 | 2.1 |

“-”, the differential expression was detected to be normal.

### GRX and TRX systems response

In yeast cytosol, it contains two prominent systems against ROS and xenobiotics, the glutaredoxin (GRX) system and the TRX system (25). In this study, at the early transcriptome profile many genes concerning those two above systems were slightly activated (Table 4 and Fig. 4). The expressions of genes encoding for glutathione synthase (*GSH1*), for glutathione glutaredoxins (*GRX1*, *GRX2* and *GRX8*), for glutathione peroxidase (*GPX1*), for glutathione transferases (*GTT1*, *GOT3* and *ECM4*) and for thioredoxins (*AHP1*, *TSA2* and *SRX1*) were up-regulated. At the late lag phase, genes (except *GRX2* and *GPX1*) in the GRX system were down-regulated, and the expression of genes in the TRX system returned to normal. Surprisingly, cytosolic catalase T-encoding gene *CTT1* displayed the consistent positive-regulation over all sampling points, expression fold changes of which could be 14.3 at the early lag phase. Furthermore, the early transcriptional response of genes (*GLO1* and *GLO2*) in the toxic methylglyoxal scavenging pathway was positively activated (Table 4 and Fig. 4).

### Transcriptional regulation and other responses

Among hundreds of DEGs identified from this lag phase transcriptome profile, several TF-encoding genes (*EDS1*, *UME6*, *MET32*, *ZAP1*, *HAP4*, *CIN5*, *COM2*, *MIG1*, *RPI1*, *YOX1*, *FKH2*, *SFL1*, *STE12* and *GAL3*) were positively regulated over all sampling points, especially the increased expression of *COM2* could be up to 22 fold changes at the early lag phase (Table 5). However, the others (*AFT2*, *CRF1*, *XBP1*, *RGM1*, *MIG3*, *MSN4* and *MBF1*) displayed just up-regulation at the early lag phase (Table 5). In contrast, the transcriptional level of *HAP1*, *PDR8*, *YRR1*, *WTM1* and *SIP4* were consistently low over all sampling points except *NRM1*, *NRG2*, *SWI5*, *ACE2* and *SUT2* (Table 5).

**Table 5.**
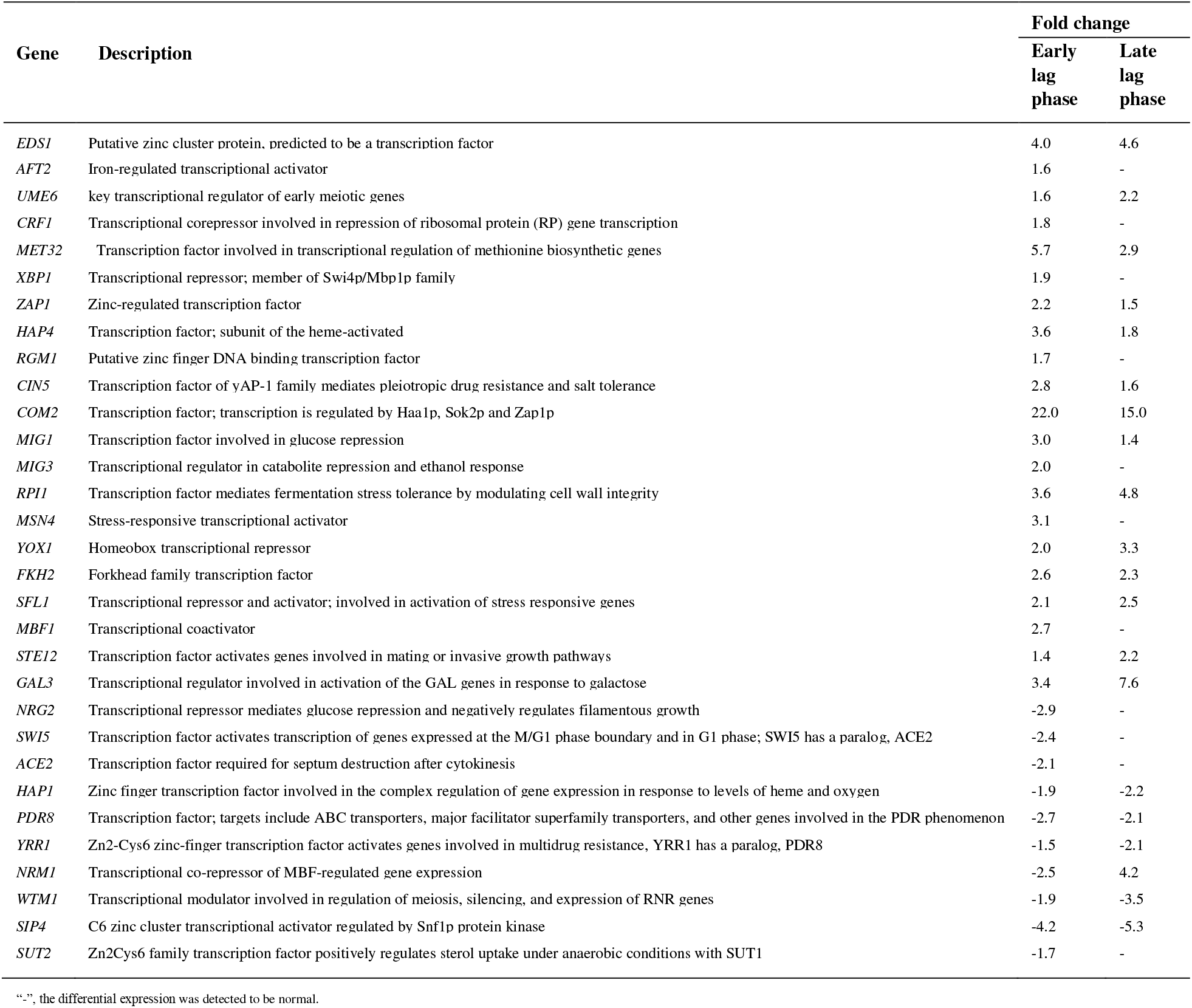
Comparative expression fold changes of TF-encoding genes responsible for the industrial *S. cerevisiae* YBA_08 response to this corn stover hydrolysate at early and late lag phases.

In addition, consistent response genes referred to ion transportation contained up-regulated genes (*PMA1* and *PMP2*) and down-regulated genes (*CTR1* and *CTR3*) (Table 3). It was noteworthy that *CTR1* and *CTR3* displayed approximately 150 and 40 fold decrease over all sampling points, respectively. Furthermore, genes in polyamine/toxin transporter-encoding genes (*TPO2*, *TPO4*, *QDR2*, *AQR1* and *ERC1*) were activated over all sampling points except *TOP3* and *DUR3* backing to normal at last (Table 3). Except that, four genes (*BNA3*, *BNA5*, *NMA1* and *PNC1*) in NAD biosynthesis and salvage, fifteen oxidoreductase-encoding genes (e.g., *AAD10*, *BDH1* and *BDH2*), and fourteen chaperone-encoding genes (e.g., *SSA1*, *SSA2*, *SSA3*, *HSP104* and *HSP82*) were up-regulated at the early lag phase (Table 2 and Table S3), whereas six genes (*CLB5*, *CLN2*, *HSL1*, *SWE1*, *CLB1* and *CLB2*) in cell cycle were repressed at the early lag phase and up-regulated at the late lag phase (Table 2 and Fig. 4). And seven genes (*ADE1*, *ADE2*, *ADE13*, *ADE17*, *RNR1*, *IMD3* and *GUA1*) in purine *de novo* biosynthesis were suppressed at the early lag phase (Table 4).

## Discussion

Actually, the lignocellulose hydrolysate generated from pretreatment contains various inhibitors, which is highly toxic to biocatalysts during biofuel fermentation. And its strong toxicity is attributed to the synergistic effect of multiple lignocellulose-derived inhibitors, which is far beyond the sum of effects caused by every single inhibitor (26, 27). To cope with negative effects under stress, such as oxidative stress, osmotic stress and cell wall stress, *S. cerevisiae* has displayed a rapid transcriptional response (12, 28). In order to explore underlying molecular mechanisms in transcriptional level, an industrial *S. cerevisiae* was employed in this study for transcriptome profile analysis in time-series under the corn stover hydrolysate. As observed in this research (Fig. 1 and Fig. 2), this corn stover hydrolysate indeed induced growth arrests and accumulations of ROS, mitochondrial and ER damages in the industrial *S. cerevisiae* YBA_08 during the lag phase, revealing negative effects existed under this corn stover hydrolysate.

### MAPK signal transduction under stress

In *S. cerevisiae*, at least three MAPK-signaling pathways, such as FG pathway, HOG pathway and CWI pathway, were verified to elicit responses under lignocellulosic inhibitors through phosphorylation of MAPKs, such as Kss1p, Hog1p, Slt2p, respectively (14, 29). Unfortunately, no information of MAPKs in those pathways was obtained in this study except that other up-regulated kinase-encoding genes *SSK22*, *RCK1*, *RCK2*, *PTP2* and *YPK2* (early), which are reported to play important roles in HOG and CWI pathways (30–33). Moreover, the significant activation of *MSN4* (early) and *STE12* (early and late) under this corn stover hydrolysate stress was found in this study, indicating Msn4p- and Ste12p-targeted genes might be activated through HOG pathway and FG pathway under this stress.(34, 35). Indeed, many target genes regulated by Msn4p and Ste12p were revealed in this research (36, 37). Although no expression information about TF-encoding genes activated by CWI pathway in this study, a key plasma membrane sensor-encoding gene *MTL1* (38) in the CWI pathway was significantly up-regulated upon this stress. Taken together, the enhanced expression of genes associated with three MAPK-signaling pathways suggested their crucial roles in the industrial *S. cerevisiae* YBA_08 adaptive response to this corn stover hydrolysate stress. Additionally, stimulations like osmotic stress may cause transient G1/G2 arrest through MAPK-mediated regulations (39, 40). Actually, our results revealed the low expression of genes (e.g., *CLB5*, *CLN2*, *CLB1* and *CLB2*) essential for Hog1p-mediated regulation of cell cycle (Fig. 4) (39, 40), together with the up-regulation of *FUS3* associated with G1 arrest (41), indicating that the industrial *S. cerevisiae* YBA_08 might display a cell cycle arrest at the early lag phase under this corn stover hydrolysate stress. Overall, the further investigation of the relevance of three MAPK-signaling pathways to this stress need to be done later.

### Adaptive stress responses

#### Oxidative stress response

Application of lignocellulose-derived inhibitors, such as furfural, HMF, acetic acid and vanillin, to *S. cerevisiae* can induce accumulations of ROS, which truly does oxidative damages to cellular compounds, such as DNA, proteins and lipids (42). This ROS accumulation result (Fig. 2) of the early lag phase obtained in this study was consistent with the above report, demonstrating that the industrial *S. cerevisiae* YBA_08 might transiently activate cellular ROS scavenging systems like catalases, GRX system and TRX system against oxidative stress (25). Indeed, this observation of early up-regulation of genes (*GSH1*, *GRX1*, *GRX2*, *GPX1*, *GOT3*, *ECM4*, *AHP1*, *TSA2*, *SRX1*, *TRX3*, *PRX1* and *CTT1*) involved in GRX system, TRX system and hydrogen peroxide detoxification under this corn stover hydrolysate (Fig. 4), greatly confirmed the above assumption. In *S. cerevisiae* GRX system, ROS-mediated damages and xenobiotics can be removed by converting the reduced form (GSH) to the oxidized disulfide form (GSSG) (43), and Gsh1p (a glutathione synthase) is an essential enzyme during oxidants detoxification (44, 45). Moreover, the efficiency of *S. cerevisiae* GRX system primarily depends on cooperations of three major parts, glutaredoxins (GRXs), glutathione peroxidases (GPXs) and glutathione transferases (GSTs) (43). *GRX1* and *GRX2*, classic GRXs-encoding genes, are regulated by STRE upon oxidative stress (46). The GPXs-encoding gene *GPX1* can protect membrane lipids against peroxidation through acting as hydroperoxides (47). And these GSTs-encoding genes *GTT1*, *GTO3* and *ECM4* possess major roles in ROS and xenobiotics detoxification through glutathione conjugate pumps (48–50). Additionally, *S. cerevisiae* contains two independent cytoplasmic and mitochondrial TRX systems in response to oxidative stress. The main components of these systems including cytoplasmic peroxiredoxins (Tsa2p, Ahp1p and Srx1p), mitochondrial peroxiredoxin Prx1p, mitochondrial thioredoxin (Trx3p and Grx2p), act as antioxidants and exhibit great functions in ROS scavenging (25). In summary, these results indeed revealed the early oxidative defense mechanism in cytosol and mitochondria under this corn stover hydrolysate stress. Further, the subsequent experiments should be carried out to explore the correlation between the key metabolite content in these systems and this stress resistance.

#### Osmotic stress response

To resist kinds of stresses like oxidative stress and osmotic stress, yeast can reprogram cellular metabolism pathways to accumulate protectants, such as trehalose, glycogen and glycerol, to preserve the integrity of plasma membrane (51). Based on this transcriptome profile, not only were several key genes (*UGP1*, *TPS1*, *TPS2* and *GSY2*) in trehalose and glycogen biosynthesis activated at the early lag phase, but degradation-related genes (*NTH1*, *NTH2*, *GDB1* and *GPH1*) were also positively regulated (Fig. 4). This ineffectual cycling of trehalose and glycogen metabolism is also observed in other research (52), probably indicating the rational distribution between energy supplies and protectant demands under this corn stover hydrolysate stress where cellular metabolisms were restricted.

Additionally, as the major solute for osmoregulation in yeast, import and export of glycerol depend on specific channels like the Fps1p-/Stl1p-facilitated channel because of the low permeability in membrane (53). Upon hyperosmotic shock, the Hog1p-mediated phosphorylation leads to removal of Ask10p (a positive regulator of Fpslp) from Fpslp and making the channel closed to prevent loss of internal glycerol, while water efflux is mediated for cell volume maintenance through suppression of *AQY2* and activation of *AQY3* (54). Also, the glycerol-H^+^ symporter Stl1p can be activated by Hog1p to import glycerol from surrounding under hyperosmotic stress (54). And Gup2p is reported to be responsible for glycerol uptake and can be activated under osmotic stress (55). Fortunately, except for *FPS1*, this early transcriptome profiling of all above mentioned glycerol transportation-related genes (Table 1 and Fig. 4) under this corn stover hydrolysate stress was successfully obtained, including up-regulation of *ASK10*, *AQY3*, *STL1* and *GUP2*, and down-regulation of *AQY2*. Meanwhile, we also observed up-regulation of glycerol metabolism-related genes (e.g., *GPP1* and *DAK2*) (Fig. 4) at the early lag phase, which was consistent with previous omics profiling results of yeast under stress like ethanol and lignocellulosic inhibitors (52, 56). Taken together, it might be reliable to speculate that the industrial *S. cerevisiae* YBA_08 was encountered with a hyperosmotic shock at the early lag phase under this corn stover hydrolysate stress. And expression differences of glycerol metabolism-related and transportation-related genes between previous reports and this observation might be attributed to the fact that industrial yeasts are often selected to ensure their ethanol fermentation efficiency due to their better responses than laboratory yeasts upon this complex stress condition (57).

#### Structural response

Rearrange of plasma membrane compositions (e.g., unsaturated fatty acid and ergosterol) to maintain its integrity has been a determinant for tolerance of stress in yeast (42). Up-regulations of unsaturated fatty acids (UFAs) metabolism-related genes and ergosterol biosynthesis-related genes are found under acetic acid, furfural and ethanol (12, 52). Surprisingly, this early lag phase investigation of a transcription decrease of essential UFAs metabolism-related genes (e.g., *FAS1* and *OLE1*) and most of ergosterol biosynthesis-related genes (e.g., *ERG10*, *ERG12* and *ERG5*) is completely in contrast to the above result (Fig. 4). As reported, upon stress the membrane integrity can be kept not only by accumulations of UFAs and ergosterol but also by enhancements of osmolytes like trehalose, which still plays an important role in stabilizing membrane proteins (58). Together with up-regulation of trehalose metabolism-related genes obtained in this study, it was rational to come to a conclusion that the industrial *S. cerevisiae* YBA_08 might prevent cell membrane from this corn stover hydrolysate damage in other ways at the early lag phase. And it might benefit for supplying of sufficient cofactor NADPH for detoxification of inhibitors like HMF, furfural and vanillin through restraining NADPH consumption in NADPH-dependent fatty acid metabolism (23, 59). Alternatively, the transient interruption of fatty acid metabolism might be a result of the feedback of the disorder of ER membrane (Fig. 2) observed in this study (23). With regard to the early transient repression of ergosterol biosynthesis in this study, it appeared to be correlated to enhancements of SAM biosynthesis and HOG pathway (60, 61). And it might be enhanced due to the late activation of three genes (*ERG3*, *ERG11* and *ERG5*) under this stress. Accordingly, plasma membrane can be disrupted by toxins like acetic acid, leading to increase of membrane permeability and eventual cytosolic acidification. To cope with it, yeast often activates H^+^-ATPase to pump out proton (62). This observation of up-regulation proton extrusion-related genes (*PMA1*, *PMP2* and *HSP30*) was consistent with the opinion (Table 3). Moreover, the significant down-regulation of two plasma membrane Cu^2+^ transporter-encoding genes (*CTR1* and *CTR3*) (63) also suggested the change in plasma membrane permeability. Besides that, other structural responses to this corn stover hydrolysate stress were exhibited in this research, including peroxisome responses and mitochondria responses. Peroxisomal fatty acid β-oxidation in yeast requires Pxa1p (peroxisomal ABC transport), Pox1p (fatty-acyl coenzyme A oxidase), Pot1p (3-ketoacyl-CoA thiolase) and Fox2p (3-hydroxyacyl-CoA dehydrogenase) (24). The altered expression of these above genes (Table 2 and Fig. 4) displayed in this study might suggest a block in peroxisomal fatty acid degradation under this stress. Reportedly, mitochondria play a crucial role in various cellular activities (energy production and general metabolism) in yeast (64) and their disorder can be found upon lignocellulose inhibitors (28, 65). Consequently, this experimental evidence of mitochondrial damages (Fig. 2) under the corn stover hydrolysate stress, together with repressions of key genes (Table 3 and Fig. 4) in mitochondrial metabolism, suggested mitochondria were severely damaged and their metabolisms were suppressed under this stress. Also, mitochondria are the main source of ROS, which can result in their dysfunction (66). In order to depose the risk of dysfunction mitochondria, mitophagy is promoted to participate in mitochondria removal, which depends on key factors Atg32p, Atg8p and Atg33p (67). Interestingly, transcription abundances of *ATG8*, *ATG32* and *ATG33* in this study were slightly high (Table 3 and Fig. 4), indicating that the industrial *S. cerevisiae* YBA_08 might initiate mitophagy to remove dysfunction mitochondria under this stress at the early lag phase.

#### Heat shock proteins (HSPs) chaperone

Stress conditions like ethanol and heat stress have been verified to lead to proteins denaturation and dysfunction (25). Against this effect, HSPs are induced to assist in chaperoning newly translated proteins, refolding misfolded proteins, and degrading aggregated proteins, and activations of HSPs are usually controlled by both Hsf1p (a heat shock TF) and Msn2p/Msn4p (25). In agreement, up-regulations of Hsps-encoding genes (Table S3 and Fig. 4) in this study suggested Hsps were also critical for tolerance of this corn stover hydrolysate stress. Ssa1p, Ssa2p, Ssa3p and Sse2p, as members of Hsp70 family, can protect nascent polypeptides and assist in refolding damaged proteins (68). Hsp82 and Cpr6p, the Hsp90 chaperone and cochaperone, are required for refolding specific target proteins and interacting with unfolded protein to assist in maturation (69). Unlike those above Hsps, Hsp104 (a member of Hsp100 family) is capable of mediating misfolded proteins to disaggregate through Hsp104/Hsp70 cooperation (25), whereas two sHSPs (small HSPs) Hsp26 and Hsp42 appear to bind unfolded proteins to promote protein solubility under stress (70). In addition, two Hsp40/J chaperones, Sis1p and Apj1p, play modulatory roles in substrate interaction with Hsp70 and SUMO-mediated protein degradation, respectively (71). The membrane organization-related protein Hsp12 has a function in stabilizing membranes under stress (72). Further, abundances of reported mitochondrial chaperones-encoding genes (*ECM10* and *HSP78*) function in protein translocation and misfolded proteins disaggregation (68) also displayed up-regulations under this stress.

#### Other responses

Recently, a report has revealed roles of *ADE1*, *ADE13* and *ADE17* associated with purine *de novo* biosynthesis in tolerance of stimuli like acetic acid and ethanol, and their transcription abundances can be enhanced after adding zinc ion upon acetic acid (73). Conversely, we found that several genes (e.g., *ADE1*, *ADE17* and *IMD3*) in purine *de novo* biosynthesis (Table 4 and Fig. 4) showed significant decreases in expression level under this corn stover hydrolysate. The zinc homeostasis-related TF-encoding gene *ZAP1* (74) was significantly up-regulated in this research. Based on these results, we speculated that upon this stress Zap1p was induced to maintain cellular suitable concentration of zinc ion for resistance, and the repression of purine *de novo* biosynthesis-related genes might be an adjoint response to this zinc ion concentration. Fantastically, most of genes in sulfur metabolism and methionine uptake (Table 4 and Fig. 4) showed high expression abundances under this corn stover hydrolysate stress at the early lag phase, appearing possible that SAM biosynthesis was enhanced. SAM is reported to be essential for the synthesis of polyamines (75). Spermidine, a member of Polyamines, can enhance yeast tolerance against fermentation inhibitors through exogenous addition or overexpression of its biosynthesis-related genes (76), together with evidences that spermine and spermidine can mediate protection against oxidative stress and osmotic stress (75, 77), suggesting that polyamine might act a role in assisting the industrial *S. cerevisiae* YBA_08 for resisting to this stress. Lucklessly, the transcription profile of genes in polyamine biosynthesis like *SPE1*, *SPE2*, *SPE3* and *SPE4* was not obtained in this study, except for up-regulations of polyamine transporter-encoding genes including *TPO2*, *TPO3* and *TPO4* (Table 4 and Fig. 4), which are important for polyamines uptake and excretion (78, 79). Also, SAM is a precursor of GSH (80), which play a major role in GRX system for removal of ROS-mediated damages and xenobiotics (25). Nevertheless, the further investigation whether this corn stover hydrolysate can result in accumulations of polyamine or SAM should be made. As reported, some oxidoreductase-encoding genes like *ADH7*, *GRE2* and *ALD4* are involved in tolerance of lignocellulosic inhibitors (52). Consistently, we also examined out the up-regulation of oxidoreductase-encoding genes (e.g., *AAD10*, *BDH1* and *BDH2*) (Table S3 and Fig. 4), demonstrating their important roles in conferring resistance to this corn stover hydrolysate. Furthermore, exposure of lignocellulosic inhibitors can cause alteration of amino acid metabolism (15, 28). As observed, genes (Table S3) involved in glycine degradation (*GCV1*, *GCV2* and *GCV3*) displayed suppressions under this corn stover hydrolysate stress. *GCV1*, *GCV2* and *GCV3* are proved to be related to purine *de novo* biosynthesis and tetrahydrofolate metabolism (81). Finally, efficient supply of NAD(P)H for detoxification toxins like ROS, aldehyde and xenobiotic to keep redox balance can benefit for resisting lignocellulosic inhibitor stress (11), and genes (*BNA3*, *BNA5*, *NMA1* and *PNC1*) are crucial in NAD biosynthesis and salvage pathway (82), indicating enhanced expressions of those four genes upon this corn stover hydrolysate stress might be somehow correlated with it.

### Regulation of TFs under stress

As mentioned above, 31 TF-encoding genes display significant changes in expression abundances according to this transcriptome profile, including 21 up-regulated genes and 10 down-regulated genes (Table 5). Besides *MSN2* and *STE12* discussed, activated stress-related TF-encoding genes contained not only early response genes (*MIG3* and *MBF1*) but also continuous response genes (*CIN5*, *SFL1* and *RPI1*). The TF Mig3p, a zinc finger protein, is induced by DNA damages and plays a regulatory role upon ethanol stress (83). Also, DNA replication stress can enhance the transcription abundance of Mbf1p (a transcriptional coactivator) (84). As a member of YAP-1 family, Cin5p can mediate resistances upon conditions like salt stress and oxidative stress (85, 86). The TF Sfl1p, a heat shock factor-like protein, can act as an activator/repressor to regulate transcriptions of stress-responsive genes like *HSP30* and *SPI1* under ethanol and heat shock (87, 88). The TF Rpi1p can modulate cellular metabolism to tolerate fermentation stress through adjusting cell wall integrity (89). Overall, those above stress-related TFs might make a significant contribute to exit from the lag phase upon this corn stover hydrolysate.

Additionally, it reports that yeast growth and cell viability are blocked under the lignocellulosic inhibitor stress (5, 6), thus we speculated that cell cycle-regulated TFs should exhibit transcriptional changes. According to this investigation of cell cycle-regulated TFs, three down-regulated genes (*ACE2*, *SWI5* and *NRM1*) and three up-regulated genes (*XBP1*, *YOX1* and *FKH2*) were found, which might somehow confer this corn stover hydrolysate tolerance through mediating G1/G2 arrest (90–94). Interestingly, we found key meiosis-regulated TF-encoding genes *COM2* and *UME6* showed high transcription abundances under this corn stover hydrolysate stress, indicating expressions of early meiotic genes might be repressed and the spore formation of the industrial *S. cerevisiae* YBA_08 might be hindered. Namely, in presence of glucose the industrial *S. cerevisiae* YBA_08 seemed impossible to tolerate this stress by sporulation (95, 96). With regard to the regulation of adaptive metabolism under stress, *EDS1*, *GAL3* and *MET32* were reported to be involved in transcriptional regulation of glucose transporter, galactose metabolism and sulfur metabolism, respectively (97–99). In other words, it might partially explain activations of genes in three above bioprocesses observed under this corn stover hydrolysate stress. Taken together, this evidence of TF-encoding genes partially might provide an access to understanding the metabolism regulation network in the adaptive response of the industrial *S. cerevisiae* YBA_08 to this stress.

#### Conclusions

Although many efforts have been made to explore the underling mechanism of yeast resisting to single or multiple lignocellulose-derived inhibitors during fermentation and some mechanisms of adaptive tolerance have been revealed, it is still known a little about the industrial yeast response mechanism to the actual pretreated lignocellulosic hydrolysate, due to variable inhibitors and their unclear synergistic effects. In this study, the comparative transcriptomic analysis of the industrial *S. cerevisiae* YBA_08 revealed numerous genes associated with MAPK-signaling pathway, oxidative stress response, osmotic stress response, Hsps chaperone, purine metabolism, SAM metabolism and TF regulation displayed a significant alteration under this corn stover hydrolysate stress during the lag phase (Fig. 4). In our opinion, it is the first time to systematically address how an industrial yeast respond to the lignocellulosic hydrolysate, providing a molecular fundament for understanding the synergistic effect of various lignocellulosic inhibitors and the regulatory mechanism of tolerance of industrial yeasts under this stress.

## Materials and Methods

### Yeast strains culture and chemicals

The *S. cerevisiae* YBA_08 (26S rDNA sequence, see GenBank no. KF141699.1, and ITS sequence, see GenBank no. MN158119.1), a diploid industrial strain, was obtained from a liquor industry (Sichuan, China). Yeast peptone dextrose (YPD) medium containing 20 g/L peptone, 10 g/L yeast extract, and 20 g/L glucose was used for cultivation of *S. cerevisiae*. YP medium supplemented with corn stover hydrolysate (YPH) was used for subsequent RNA-seq analysis of *S. cerevisiae*. Strains were stored in 30% glycerol stock at −80 °C and cultivated for 24 hours at 30 °C and 200 rpm. Corn stover hydrolysate was prepared as described with some changes (100). Firstly, oven-dried corn stover particles, milled and screened by 40 mesh sieve, were thoroughly mixed with diluted (2% w/w) sulfuric acid as catalyst for 24 hours and incubated at 121 °C with steams for 1 hours. Secondly, the pH of acidic corn stover slurry was adjusted to the optimal range for 72 hours enzymatically hydrolyzing with cellulase. Finally, after sterilization through filter membrane (0.22 μm), the corn stover hydrolysate was completely prepared. All reagents were purchased by Sigma-Aldrich (St. Louis, MO, USA), Difco (Detroit, MI, USA), Sangon Biotech (Shanghai, China), and Best-Reagent (Chengdu, China).

### RNA-seq and bioinformatics analysis

*S. cerevisiae* YBA_08 was cultivated in YPH at 30 °C and 200 rpm. For RNA-seq samples isolation, the suspensions were harvested (8,000 rpm, 4 min, 4 °C) after incubation at different time points (3 h and 6 h), the cell pellets were flash-frozen in liquid nitrogen and stored at −80 °C before analysis. Subsequently, they were sent to Biomarker Technologies (Beijing, China) for RNA extraction, cDNA library conduction and sequencing. The RNA samples were sequenced on an Illumia HiSeq 2500 platform and data were analyzed on Biomarker Cloud (https://international.biocloud.net). The differentially expressed genes (DEGs) were identified as the significantly regulated genes by using thresholds (fold changes ≥ 2 and p-value ≤ 0.01). Functional catalogues analysis of DEGs was performed through MIPS Functional Catalogue (101).

### ROS accumulation, mitochondrial and ER membrane damages analysis

After *S. cerevisiae* YBA_08 growing in YPH at 30 °C and 200 rpm for 3 hours and 6 hours, respectively, the cell pellets were stained by 2’ 7’-dichlorofluorescein diacetate (DCF) (Sigma-35845), MitoTracker Green FM (Thermo Fisher Scientific E34250) and ER-Tracker^™^ Red dye (Thermo Fisher Scientific E34250) for ROS accumulation, mitochondrial membrane damages analysis and endoplasmic reticulum (ER) membrane damages analysis, respectively (65). Briefly, the suspension (1 mL) of yeast culture was centrifuged with 12,000 rpm to obtain the cell pellets. After staining for 0.5-2 hours at 30 °C, the cell pellets were washed with sterilized deionized water and suspended in 0.1 M phosphate buffer saline (PBS) (pH 7.0). The suspension (1 μL) with stained cell pellets was placed under GFP filter or Rhod filter lens of Axio Imager A2 for observation, and at least 100 cells should be quantified for each time. The concentrations of DCF, MitoTracker Green FM and ER-Tracker™ Red dye for staining were 2.5 mg/ml, 1mM and 1μg/μL, respectively. After adding in the suspension the final concentrations of DCF and MitoTracker Green FM were 20 μM and 20-200 nM, respectively.

## Data Availability

The RNA-seq data have been deposited in the Sequence Read Archive (SRA) of NCBI (102) and are accessible with a number SRP251210 (https://www.ncbi.nlm.nih.gov/sra/SRP251210).

## Supplemental Material

**Supplemental file 1. Fig. S1.** Transcriptome differential expression analysis for the industrial *S. cerevisiae* YBA_08 under this corn stover hydrolysate stress. **Table S1.** Functional categories of early DEGs responsible for the industrial *S. cerevisiae* YBA_08 tolerance to this corn stover hydrolysate. **Table S2.** Functional categories of late DEGs responsible for the industrial *S. cerevisiae* YBA_08 tolerance to this corn stover hydrolysate. **Table S3.** Comparative expression fold changes of other stress-related genes for the industrial *S. cerevisiae* YBA_08 response to this corn stover hydrolysate at early and late lag phases.

## Funding

This work was supported by the National Natural Science Foundation of China (No. 31570086), the 2011 Collaborative Innovation Center for Farmland Protection and Agricultural Product Safety in Sichuan Province, and the Talent Introduction Fund of Sichuan Agricultural University (No. 01426100).

## Authors’ contributions

MGM, XLK and YPG conceived and designed research. XLK, YPG and ZYZ conducted experiments. XLK, YPG, ZYZ, XDH, XBH, DFX and QL analyzed data. HYW, XL and QC contributed analytical tools. XLK and YPG wrote the manuscript. MGM revised the manuscript. All authors read and approved the manuscript.

## Conflict of interest

The authors declare that they have no competing interests.

## Ethical approval

This article does not contain any studies with human participants or animals performed by any of the authors.

